# The Effects of Unilateral Transcranial Direct Current Stimulation on Unimanual Laparoscopic Peg-Transfer Task

**DOI:** 10.1101/2021.04.13.439617

**Authors:** Zaeem Hadi, Aysha Umbreen, Muhammad Nabeel Anwar, Muhammad Samran Navid

## Abstract

**Introduction:** Efficient training methods are required for laparoscopic surgical skills training to reduce the time needed for proficiency. Transcranial direct current stimulation (tDCS) is widely used to enhance motor skill acquisition and can be used to supplement the training of laparoscopic surgical skill acquisition. The aim of this study was to investigate the effect of anodal tDCS over the primary motor cortex (M1) on the performance of a unimanual variant of the laparoscopic peg-transfer task.

**Methods:** Fifteen healthy subjects participated in this randomized, double-blinded crossover study involving an anodal tDCS and a sham tDCS intervention separated by 48 hours. On each intervention day, subjects performed a unimanual variant of laparoscopic peg-transfer task in three sessions (baseline, tDCS, post-tDCS). The tDCS session consisted of 10 minutes of offline tDCS followed by 10 minutes of online tDCS. The scores based on the task completion time and the number of errors in each session were used as a primary outcome measure. A linear mixed-effects model was used for the analysis.

**Results:** We found that the scores increased over sessions (*p*<0.001). However, we found no effects of stimulation (anodal tDCS vs. sham tDCS) and no interaction of stimulation and sessions.

**Conclusion:** This study suggests that irrespective of the type of current stimulation (anodal and sham) over M1, there was an improvement in the performance of the unimanual peg-transfer task, implying that there was motor learning over time. The results would be useful in designing efficient training paradigms and further investigating the effects of tDCS on laparoscopic peg-transfer tasks.

## 1 Introduction

Laparoscopic surgery is a minimally invasive method of surgery with advantages over conventional techniques, including a reduction in blood loss, post-operative pain, and incision scars (Klempous et al., 2018). Since laparoscopic tools are inserted through a small cavity, surgeons have reduced or altered tactile-sensation, limited degrees of freedom of movement, and indirect vision during a laparoscopic procedure (Gallagher & O’Sullivan, 2012; Spruit, Kleijweg, Band, & Hamming, 2016). With the increase in demand for minimally invasive surgical procedures, the requirement for associated technical skills (Spruit et al., 2016) and appropriate training procedures have also increased.

The set of processes encompassing skill acquisition and motor adaptation brings about a relatively permanent change in a person’s behavior and is known as motor learning (Nieuwboer, Rochester, Müncks, & Swinnen, 2009; Schmidt, Lee, Winstein, Wulf, & Zelaznik, 2018). The principles of motor learning are also applicable to learning laparoscopic skills (Spruit et al., 2016). Numerous methods are available for learning laparoscopic skills, such as training using animal models, simple box trainers, and virtual reality (VR) based trainers (Palter, Orzech, Aggarwal, Okrainec, & Grantcharov, 2010). Simple laparoscopic trainers are preferable in terms of cost-effectiveness (Nguyen, Braga, Hoogenes, & Matsumoto, 2013). Laparoscopic training tasks include peg- or bead-transfer, pattern cutting, intra-, and extra-corporeal knots, placement of a mesh over a defect, placement of a ligating loop, and placement of a clip at appropriate positions (Derossis et al., 1998).

Many studies have shown that motor learning can be enhanced by increasing cortical excitability (Boggio et al., 2006; Nitsche, Schauenburg, et al., 2003; Stagg et al., 2011). One of the commonly used methods for modulating neural plasticity is transcranial direct stimulation (tDCS) (Paulus, 2011). tDCS is a non-invasive brain stimulation technique for modulating motor cortex excitability by applying small amounts of direct current on a person’s scalp (Nitsche & Paulus, 2000). The current is applied using two or more electrodes depending upon the configuration; however, a simple bipolar configuration is composed of two electrodes called anode and cathode. Current travels through the anode into the brain tissue and returns to the cathode, stimulating the underlying cortical neurons. The electrode at the stimulation site is usually termed as an active electrode. Although some current is lost at scalp, a substantial amount of current penetrates the brain to modify the excitability levels of underlying neurons (Bolognini, Rossetti, Casati, Mancini, & Vallar, 2011; Nitsche, Liebetanz, et al., 2003; Zaghi, Acar, Hultgren, Boggio, & Fregni, 2010). These effects are reversible and can usually last up to an hour or more after stimulation and are dependent upon the duration of stimulation (Nitsche & Paulus, 2001). In terms of mechanism of action, tDCS mediates the transmembrane potentials of cortical neurons and does not act by inducing the action potentials (Bolognini et al., 2011). The modulation of sodium and calcium gated channels and N-methyl-D-aspartate (NMDA) receptor activity produces effects similar to long-term potentiation (LTP) and long-term depression (LTD), that are associated with neuronal plasticity (Liebetanz, Nitsche, Tergau, & Paulus, 2002; Nitsche et al., 2004).

Since anodal tDCS enhances cortical excitability, it is assumed that it may also improve motor learning by enhancing neuroplasticity (Reis & Fritsch, 2011). This assumption has been tested and proven correct in various studies. For example, anodal tDCS over the primary motor cortex (M1) has been found to improve motor learning in implicit (Nitsche, Schauenburg, et al., 2003) and explicit motor learning tasks (Stagg et al., 2011), visuomotor task performance (Antal et al., 2004), and performance of non-dominant hand in hand function test (Boggio et al., 2006). However, the effect of tDCS over the primary motor cortex (M1) could also be task-specific since no improvement has also been reported on bimanual motor task (Vancleef, Meesen, Swinnen, & Fujiyama, 2016).

Recently, there has been some interest in the application of tDCS for the enhancement of laparoscopic skill acquisition. When we started this study, only one study (P. Ciechanski et al., 2018) looked at the effects of tDCS on laparoscopic training and found no significant effect of tDCS on the peg-transfer task; however, a significant improvement for the pattern-cutting task was observed. Despite no significant improvement in the peg-transfer task, an appropriately powered study could detect a medium effect size. Moreover, the commonly used peg-transfer task is bimanual; thus, it is expected that the application of anodal tDCS on the dominant side alone may not improve the performance of a bimanual task (P. Ciechanski et al., 2018; Vancleef et al., 2016). Similar results for the effects of tDCS on bimanual performance improvement of peg-transfer task were found by two recent studies (Patrick Ciechanski et al., 2019; M. L. Cox et al., 2020).

Since it is possible that the non-significant findings in previous studies are due to bimanual tasks and stimulation at only the dominant hemisphere, we conducted this study to evaluate if tDCS of dominant hemisphere can affect *unimanual* tasks. Therefore, we designed a double-blinded randomized crossover study, which was aimed specifically to assess the effect of anodal tDCS over M1 on the performance of a unimanual variant of the laparoscopic peg-transfer task.

## 2 Materials and methods

The study was approved by the Institutional Ethical Committee, National University of Sciences and Technology (NUST), Islamabad, Pakistan. We carried out sample size calculations based on the effect size from a previous study (P. Ciechanski et al., 2018) using G*power version 3 (Faul, Erdfelder, Lang, & Buchner, 2007). A sample size of 16 was required to detect a Cohen’s d of 0.40 with a power of 90%.

The data for this study were collected in the experimental room of Human Systems Lab, School of Mechanical and Manufacturing Engineering, National University of Sciences and Technology, Islamabad, Pakistan, from March 2018 to August 2018.

### 2.1 Participants

Before participation, seventeen healthy young university students (10 males, 26.24±2.44 years) able to provide consent were screened based on any metallic implants, type of prescribed or unprescribed medication, pregnancy, history of seizure, history of the neurosurgical procedure, and neurological or psychiatric condition. Subjects were then tested for their hand preference using the “Dutch Handedness Questionnaire”. One subject was excluded because of left-hand preference. Sixteen right-handed subjects then signed an informed consent form and were enrolled in the study.

### 2.2 Experiment Protocol

The experiment protocol is illustrated in Figure 1. The subjects were required to visit the lab for two days for the experiment. The screening was done before the first visit or on the day of the first visit. The second visit of each subject was scheduled after 48 hours of the first visit. On the first visit, an experimenter demonstrated the use of equipment (i.e., laparoscopic graspers and the trainer) to the subjects. Afterward, the subjects had 15 minutes of practice session in which the subjects practiced transferring beads. During practice session, subjects could place beads at any place on the laparoscopic trainer and were given feedback about the errors. There was a 5 minutes break after the practice session, during which the experimenter explained the peg-transfer task to the subjects. Afterward, the subjects performed the whole peg-transfer task three times.

**Figure 1.**
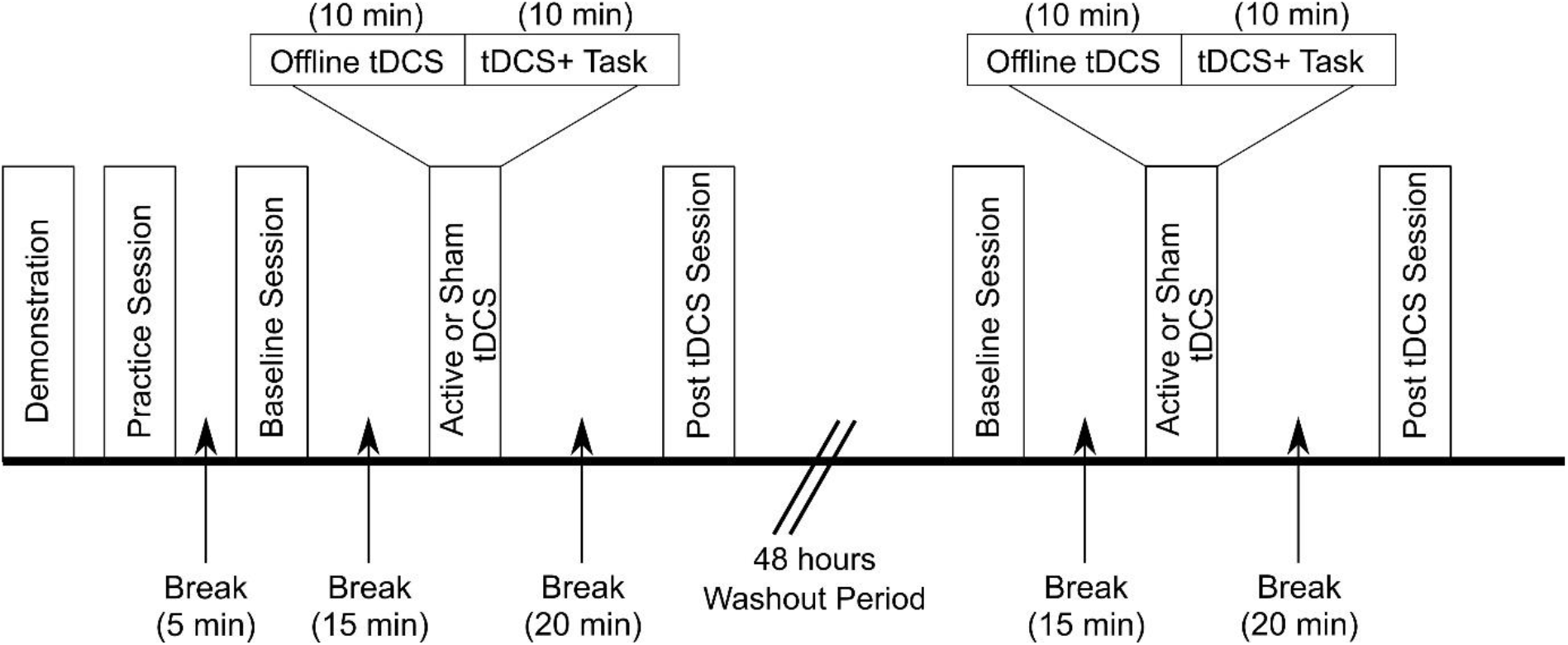
Experiment Protocol.

On each visit, the subjects participated in three experimental sessions: baseline, online-tDCS, and post-tDCS. The subjects performed the peg-transfer task in each session. There was a break of 15 minutes between the baseline and online-tDCS sessions and a break of 20 minutes between the online-tDCS and post-tDCS sessions. Subjects were given anodal or sham stimulation in the online-tDCS session. The order of stimulation was randomized through a simple random number generator for all subjects. In the online-tDCS session, stimulation started 10 minutes prior to the beginning of the task and continued for another 10 minutes during the task. The task lasted for approximately 10 minutes; therefore, the stimulation was given for a total of 20 minutes. Videos of the task performance were also recorded.

The study was double-blinded; therefore, the experimenter and the subjects were blinded to the stimulation condition. An independent researcher handled the device during the stimulation sessions.

### 2.3 Laparoscopic Trainer

We used a custom-built digital laparoscopic trainer based on object detection sensors. The position of each bead placement was detected digitally, and the timing information of individual bead placement and task start and completion time were stored. The trainer had a time resolution of 1 millisecond and a total of 144 locations for bead placement to allow the customization of pattern making. The trainer was placed inside a stage with adjustable height and lightning. The stage had two areas at the top for inserting laparoscopic graspers. An autofocus HD video camera was also mounted at the top of the stage. The video was visible to the subject on a screen placed approximately 1.5 meters away from the subject. The subjects used a standard, double action, universal atraumatic laparoscopic grasper of diameter 5 mm and length 31 cm from GERATI Healthcare Pvt. Ltd. Pakistan, for performing the peg-transfer task.

### 2.4 Peg-Transfer Task

To assess the unimanual performance enhancement, we used a modified version of the peg-transfer task. The subjects were required to place beads over a pre-marked pattern (Figure 2) as quickly as possible in the three experimental sessions. The order of bead placement (Figure 2) was displayed beside the laparoscopic trainer printed on a paper. The subjects were instructed to place the beads in an anticlockwise manner, moving from location 1 to location 13 in ascending order within the marked square on the trainer board, as shown in Figure 2. The beads were available in a small container built on top of the trainer board. The subjects were required to use their right hand for holding and using the grasper. The subjects used their left hand was to orient the face of the laparoscopic grasper.

**Figure 2.**
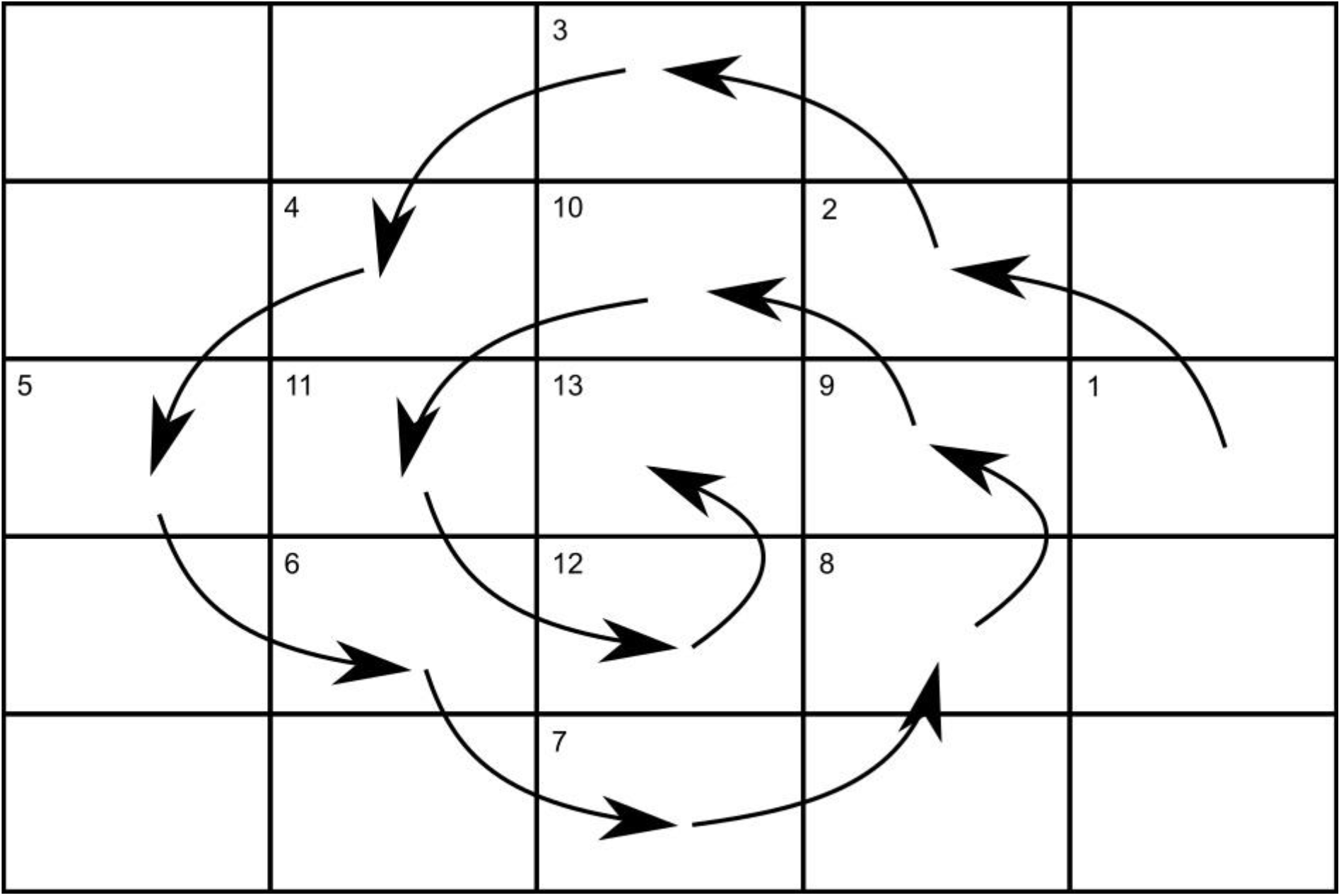
Peg-transfer task. Subjects were required to start placing beads from position 1 to position 13 in an anticlockwise manner, as depicted by arrows.

Subjects were instructed to place beads in horizontal orientation only and not touch or displace other beads while placing a new bead or while retracting the grasper. The subjects were verbally reminded of these instructions as well as the order of placement before every session.

### 2.5 tDCS

We used Caputron, ActivaTrek’s “ActivaDose 2” tDCS device for stimulation. The electrodes used as anode and cathode had a size of 3×3 cm (9 cm^2^). The current intensity was 1 mA for stimulation, and thus the current density was 0.1 mA/cm^2^. Anode was placed over left M1 (corresponding to electrode location C3 according to 10-20 standard of electroencephalography (EEG) electrode system), and cathode was placed over the right supraorbital region (corresponding to electrode location Fp2 in 10-20 standard of EEG electrode system). Sponges of the same size were inserted inside the electrodes. Sponges were soaked in the saline solution before inserting into electrodes. Electrodes were held in place with “Caputron Universal Strap”.

During anodal stimulation, the current was ramped up over 10 seconds, held constant at 1 mA for 20 minutes, and then ramped down over 3 seconds. In sham stimulation, the current was ramped up to 1 mA over 10 seconds, held constant for 1 minute, and then ramped down over 3 seconds.

### 2.6 Outcomes

Scoring metrics for laparoscopic training tasks vary depending upon the trainer used for practice and the modality of importance (i.e., task completion time, errors, or the combination of both). Usually, task completion time is used as a score after penalizing it for the errors (i.e., Score = Cut-off time – (Task completion time + (penalty * factor)) (Berg et al., 2015; Derossis et al., 1998; Matzke, Ziegler, Martin, Crawford, & Sutton, 2017). Cut-off time is pre-selected and is often 300 s in peg-transfer task, which is an approximate measure of task completion time used in Fundamentals of Laparoscopic Surgery (Derossis et al., 1998). Different types of penalty factors have also been used previously, such as fixed factor for all error types (Berg et al., 2015; D. R. Cox, Zeng, Frisella, & Brunt, 2011; Matzke et al., 2017) or different factors depending on the type of error (M. L. Cox et al., 2020). We used the same general formula to calculate the score, which we also used as our primary outcome measure. The cut-off time was selected as 600 s since our modified peg-transfer task required ten minutes approximately. The weighting factors were decided based upon the importance of the type of error. The formula for the score is given as:

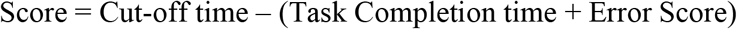

Cut-off time was 600 s and Error Score was calculated as:

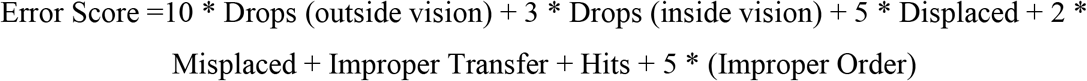

where

Drops (outside vision): Beads dropped outside the field of vision of the camera

Drops (inside vision): Beads dropped inside the field of vision of the camera

Displaced: Already placed beads displaced from their position

Misplaced: Beads placed at an unmarked location

Improper Transfer: Beads placed in inaccurate orientation or transferred in an incorrect manner

Hits: Nearby beads hit while placing a bead or retracting the grasper

Improper Order: Beads not placed in the order specified

A score of zero was awarded if the score was negative, as reported in (Korndorffer, Bellows, Tekian, Harris, & Downing, 2012) for the laparoscopic suturing task. Task completion time was available from the logged files of the laparoscopic trainer and video recordings of task performance of all subjects. A researcher, blinded to sham or active conditions, analyzed the video recordings for assessing the errors.

### 2.7 Statistical Analysis

The score was used as the primary outcome measure, as discussed in the previous section. We used the linear mixed-effects model (LMM) for analysis in IBM SPSS Statistics version 26. The subjects were specified as a random effect, whereas “Session” (baseline, tDCS, post tDCS) and “Stimulation” (Active tDCS, Sham tDCS) were specified as fixed effects. Model fit was evaluated for covariance structures of first-order autoregressive, compound symmetry, and unstructured covariance between the repeated measures using Akaike’s information criterion (AIC). The covariance structure of scaled identity was chosen for a random effect since it assumes no correlation between each subject. The first-order autoregressive covariance structure was selected since it had the lowest AIC. The main effects of Session, main effects of Stimulation, and interaction of Session and Stimulation were calculated. Results were considered significant if p<0.05, and multiple comparison correction for post-hoc results was done using Bonferroni correction where required. The data are presented as mean ± SD unless otherwise indicated.

## 3 Results

One of the 16 subjects was uncomfortable with the tDCS, and thus, the stimulation session was discontinued at 10 minutes (pre-task). We excluded the subject from the analysis, and results from 15 subjects are reported here.

The scores of subjects from all sessions are presented in Figure 3. Although there seems to be a slight difference in the subjects’ mean scores in both stimulation types, the variance was relatively high.

**Figure 3.**
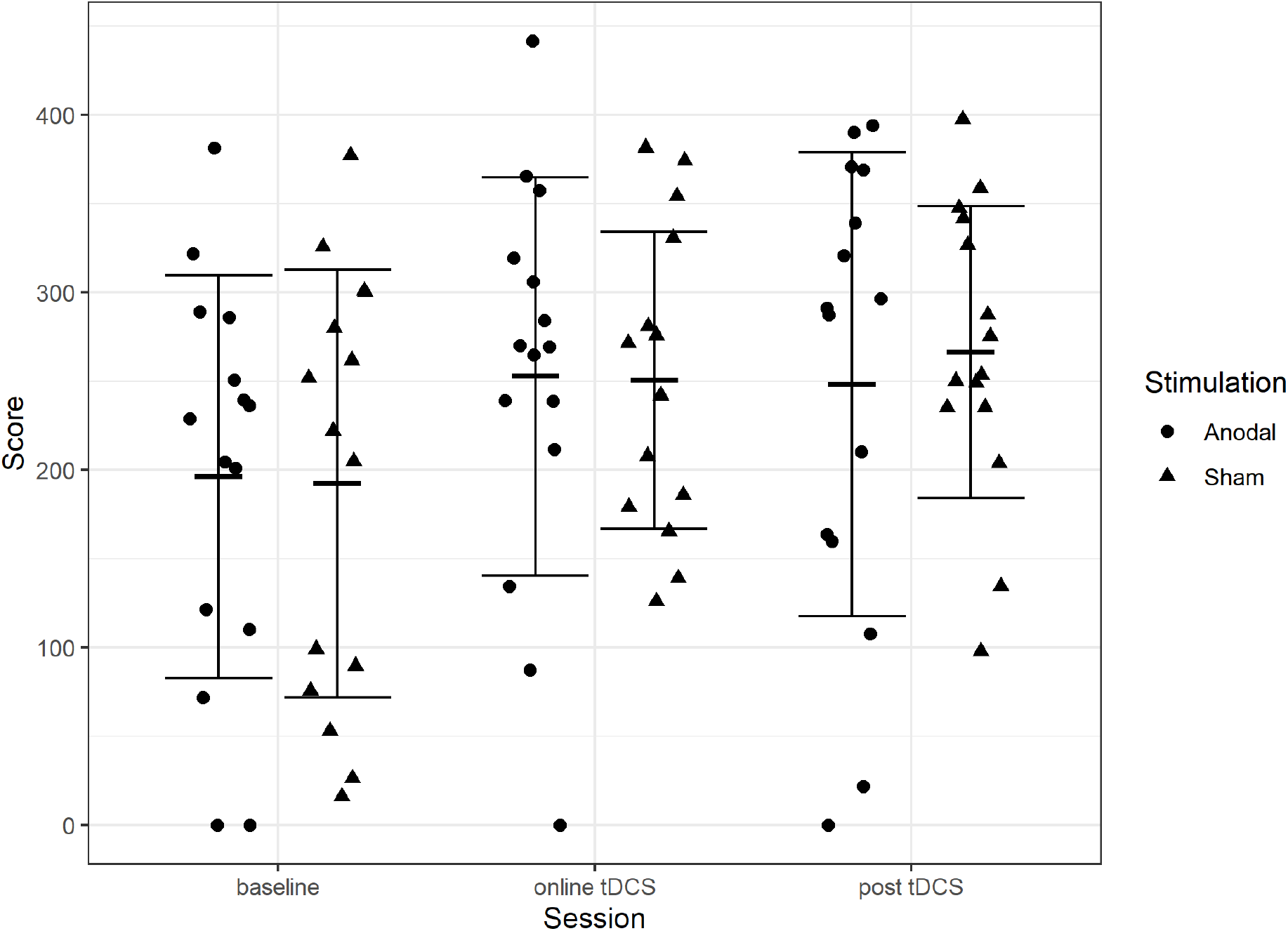
Scores. Circles and triangles show individual scores. Error bars show mean ± SD scores.

The main effects of the factors “Stimulation” and “Session” and their interaction are provided in Table 1. We found no significant interaction of factors “Stimulation” and “Session” (F(2, 55.40) = 0.19, p = 0.83), and no main effect of “Stimulation” (F(1, 19.03) = 0.03, p = 0.856). A significant effect of “Session” was observed (F(2, 58.22) = 7.76, p = 0.001). The significant effect of “Session” showed that the performance of subjects improved over time, and there was no difference in their scores because of the type of Stimulation.

**Table 1.** Type III Tests of Fixed Effects.

| Source | Numerator df | Denominator df | F | Sig. |
| --- | --- | --- | --- | --- |
| Intercept | 1 | 14.44 | 127.53 | .000 |
| Stimulation | 1 | 19.03 | .03 | .856 |
| Session | 2 | 58.22 | 7.76 | .001 |
| Stimulation * Session | 2 | 55.40 | .19 | .830 |

Estimated marginal means of the main effects of Stimulation and Session and interaction of Stimulation and Session are presented in Table 2, Table 3, and Table 4 respectively.

**Table 2.**
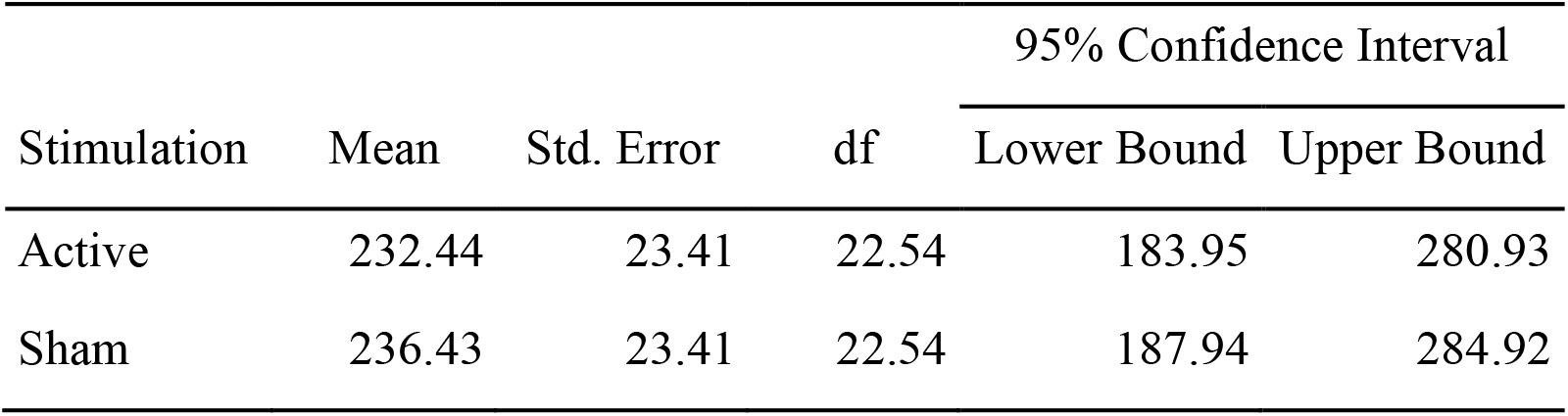
Estimated marginal means of the factor Stimulation.

| Stimulation | Mean | Std. Error | df | 95% Confidence Interval |  |
| --- | --- | --- | --- | --- | --- |
|  |  |  |  | Lower Bound | Upper Bound |
| Active | 232.44 | 23.41 | 22.54 | 183.95 | 280.93 |
| Sham | 236.43 | 23.41 | 22.54 | 187.94 | 284.92 |

**Table 3.** Estimated marginal means of the factor Session.

| Session | Mean | Std. Error | df | 95% Confidence Interval |  |
| --- | --- | --- | --- | --- | --- |
|  |  |  |  | Lower Bound | Upper Bound |
| Baseline | 194.32 | 22.96 | 21.29 | 146.62 | 242.02 |
| Online tDCS | 251.65 | 22.96 | 21.29 | 203.95 | 299.35 |
| Post tDCS | 257.34 | 22.96 | 21.29 | 209.64 | 305.04 |

**Table 4.** Estimated marginal means of the interaction Stimulation and Session.

| Stimulation | Session | Mean Score | Std. Error | df | 95% Confidence Interval |  |
| --- | --- | --- | --- | --- | --- | --- |
|  |  |  |  |  | Lower Bound | Upper Bound |
| Active | Baseline | 196.27 | 27.65 | 39.59 | 140.38 | 252.16 |
|  | online-tDCS | 252.67 | 27.65 | 39.59 | 196.87 | 308.66 |
|  | post-tDCS | 248.27 | 27.65 | 39.59 | 192.38 | 304.17 |
| Sham | Baseline | 183.446 | 27.65 | 39.59 | 136.47 | 248.26 |
|  | online-tDCS | 246.235 | 27.65 | 39.59 | 194.64 | 306.43 |
|  | post-tDCS | 269.225 | 27.65 | 39.59 | 210.51 | 322.29 |

## 4 Discussion

This study presents the effects of anodal tDCS over the M1 region on the unimanual variant of the laparoscopic peg-transfer task. We used a modified laparoscopic peg-transfer task to test unimanual performance enhancement. Our results show that there was an improvement in performance over sessions. However, this was not related to tDCS as a single session of anodal tDCS over the M1 region did not affect the laparoscopic peg-transfer task compared to sham stimulation.

It has been reported previously that unilateral tDCS does not affect the bimanual peg-transfer task (P. Ciechanski et al., 2018; Patrick Ciechanski et al., 2019) while bilateral tDCS only has an online effect on the bimanual peg-transfer task (M. L. Cox et al., 2020). Our study was focused on unimanual peg-transfer task and unilateral tDCS M1 stimulation. We found no effect of stimulation, online or offline, and thus were unable to replicate the results of the study by Cox et al. (M. L. Cox et al., 2020). This could be because we only gave stimulation in a single training session compared to other studies, which provided stimulation across multiple training sessions and possibly minimized learning over time (P. Ciechanski et al., 2018; Patrick Ciechanski et al., 2019; M. L. Cox et al., 2020). We also found no interaction effect of stimulation-type and sessions, which means that there was no post-stimulation difference in scores between active and sham stimulations. Our results suggest that unilateral tDCS may not have a significant effect on the unimanual peg-transfer task.

Although no statistically significant effects were observed for tDCS on the peg-transfer task, the discussion on effect size may provide some further explanation. Our study design was powered to detect an effect size of 0.4 with the power of 90%. This was based on (P. Ciechanski et al., 2018) which had an effect size of Hedge’s g = 0.40. Since we could not detect that effect size with an appropriately powered study, this could mean that the actual effect size is much lower than the estimated effect size. Another important thing to notice is that two of the three reported studies investigating the effect of tDCS on peg-transfer task had a negative effect size (Hedge’s g = −0.44 (Patrick Ciechanski et al., 2019), Glass’s Δ = −0.56 (M. L. Cox et al., 2020)). Thus, it is not yet clear whether tDCS improves the performance of the peg-transfer task or deteriorates it; therefore, further investigation is required to avoid using intervention with negative effects.

The peg-transfer task used in this study was modified for two reasons. One reason was to digitally measure the task timings instead of using commonly used video recordings. The second was to test for the possibility of unimanual performance enhancement. Moreover, the task we designed was relatively much difficult than the standard peg-transfer task mainly used in FLS (Derossis et al., 1998). The original peg-transfer task is a bimanual task that requires the transfer of objects from left pegs to right pegs on board and then from right to left. The objects to be transferred are larger and easier to carry or move. Thus, it would be suitable to use alternatives with different complexity levels of the task to allow further improvement and not limit performance. Yet, our task showed a main effect of sessions, which means that subjects were successfully able to improve their performance in just three sessions despite the difficulty of the task.

In terms of study design, there is evidence that motor performance can be enhanced by a single session of tDCS (Antal et al., 2004; Boggio et al., 2006; Nitsche, Schauenburg, et al., 2003). However, multi-session tDCS spanned across multiple days has been found to have much larger performance enhancements in motor skill acquisition, which is mainly attributed to the consolidation effect and not within-session learning (Reis et al., 2009). This superiority of performance can last up to months as well (Reis et al., 2009). Our study could not find an effect on performance with a single session of tDCS. Since there is a main effect of sessions, it could mean that learning is too plastic and has a large margin of improvement over sessions. Thus, it would be difficult to single out the effect of the intervention in a single session in such a scenario. The difference, if any, would be much clear if evaluated at asymptotic learning levels.

Previous literature on motor learning suggests positive effects of online tDCS on online motor learning, but the same is not established for offline learning (post-tDCS) effects (Reis & Fritsch, 2011). Offline tDCS is reported to have negative effects when applied before task performance (Stagg et al., 2011). In contrast, offline tDCS applied after the task has been reported to have no impact on offline motor learning (Chen et al., 2020). Our study used a combination of both, online tDCS and offline tDCS. However, it is unclear whether offline tDCS application had any kind of deteriorating effect as reported in (Stagg et al., 2011) since the mean score in active tDCS appears to be higher than sham tDCS.

## 5 Limitations

This study has some limitations. Considering that this is one of the earlier studies evaluating the effect of tDCS on laparoscopic peg-transfer task and the first study to use a unimanual peg-transfer task, the sample size estimates might not be accurate. We powered our study based on effect size from (P. Ciechanski et al., 2018) but failed to detect any difference in the intervention. This means a much bigger sample size is required to get an accurate estimate of true effect size. The modified peg-transfer task that we used is relatively different from the simpler versions available, considering that it was designed to accommodate the digital nature of the laparoscopic trainer. This reduces the equivalence of comparison of raw scores between studies; however, standardized comparisons would still be possible. Based on our results, we suggest that single session tDCS design may not be useful at picking up differences since the performance is much plastic and variance is high in the early stages of learning. Thus, multiple training sessions with concurrent tDCS or tDCS after asymptotic learning would reduce variance in scores and result in more accurate predictions.

## 6 Conclusion

This study suggests that single-session anodal tDCS over M1 does not significantly improve the learning of a unimanual peg-transfer task; however, these findings are not evidence of a null or negative effect. The findings would facilitate in designing more efficient training paradigms in the future for surgical residents.

## Funding

We received no formal funding for this study.

## Conflict of interest

The authors declare that the research was conducted in the absence of any commercial or financial relationships that could be construed as a potential conflict of interest.

## Author contributions

Conceptualization: AU, MNA, MSN; Data curation: ZH, AU; Formal Analysis: ZH, AU; Investigation: ZH, AU; Methodology: ZH, AU, MSN; Project administration: ZH, AU, MNA; Resources: MNA, MSN; Software: ZH, AU; Supervision: MNA, MSN; Validation: ZH, MSN; Visualization: ZH; Writing – original draft: ZH; Writing – review & editing: ZH, AU, MNA, MSN.

## Notes

### Competing Interest Statement

The authors have declared no competing interest.

